# Four-dimensional quantitative analysis of cell plate development using lattice light sheet microscopy identifies robust transition points between growth phases

**DOI:** 10.1101/2023.10.03.560767

**Authors:** Rosalie Sinclair, Minmin Wang, Zaki Jawaid, Jesse Aaron, Blair Rossetti, Eric Wait, Kent McDonald, Daniel Cox, John Heddleston, Thomas Wilkop, Georgia Drakakaki

## Abstract

Cell plate formation during cytokinesis entails multiple stages occurring concurrently and requiring orchestrated vesicle delivery, membrane remodeling, and timely polysaccharide deposition, such as callose. Such a dynamic process requires dissection in time and space; hence this has been a major hurdle in studying cytokinesis. Using lattice light sheet microscopy (LLSM) we studied cell plate development in four dimensions, monitored by the behavior of the cytokinesis specific GTPase RABA2a.

We monitored the entire length of cell plate development, from its first emergence, with the aid of RABA2a, both in the presence and absence of cytokinetic callose. By developing a robust cytokinetic vesicle volume analysis, we identified distinct behavioral patterns allowing for the identification of three easily trackable, cell plate developmental phases. Notably, the phase transition between phase I and phase II is striking, indicating a switch from membrane accumulation to the recycling of excess membrane material.

We interrogated the role of callose using pharmacological inhibition with LLSM and electron microscopy. Loss of callose inhibited phase transition, establishing quantitatively the critical role and timing of the polysaccharide in cell plate expansion and maturation.

This study exemplifies the power of LLSM, combined with quantitative analysis to decode and untangle such a complex process.

**Highlight:** We employed lattice light sheet 4D microscopy in plants to dissect cytokinesis, a multistage process involving orchestrating delivery of membranes and timely polysaccharide deposition. Robust quantitative analysis revealed distinct phase shifts, while inhibition of callose deposition abolished the phase transition.

## Introduction

Cytokinesis is a fundamental process in life, determining growth, development, and differentiation. In plant cytokinesis, a process fundamentally different from cytokinesis in animals and fungi (Samuels et al., 1995; Staehelin and Hepler, 1996; Jurgens, 2005; Drakakaki, 2015; Gu and Rasmussen, 2022; Sinclair et al., 2022), *de novo* formation of a cell plate partitions the cytoplasm of the dividing cell. Cell plate development occurs in multiple stages (Samuels et al., 1995). It requires the directed and choreographed accumulation of post-Golgi vesicles via the phragmoplast (a structure composed of cytoskeletal polymers, associated proteins, and membranes) at the division plane.

Cell plate expansion is centrifugal, led by the accumulation and fusion of vesicles arriving at the leading edge. Morphologically determined, various cell plate developmental stages exist simultaneously at cytokinesis (Samuels et al., 1995; Segui-Simarro et al., 2004). First, during the fusion of vesicle stage (FVS), cytokinetic vesicles guided by the phragmoplast arrive at the division plane where fusion occurs. Upon vesicle fusion and fission, tabulation, primarily due to the activity of dynamin-related proteins (Otegui et al., 2001; Segui-Simarro et al., 2004), helps the transition to a tubular-vesicular network (TVN) (Samuels et al., 1995; Segui-Simarro et al., 2004). The membrane morphology then evolves through the expansion of the network to a smoother tubular network (TN). The polysaccharide callose is predominantly deposited at this stage (Samuels et al., 1995). The excess membrane material is recycled via clathrin-coated vehicles (Otegui et al., 2001). As the cell plate continues to smoothen and expand, it transitions to a planar fenestrated sheet (PFS). The cell plate stiffens with the deposition of polysaccharides leading to the formation of a cross wall. As the phragmoplast expands centrifugally, delivering vesicles at the leading edge of the cell plate, the center matures with the removal of excess membranes and deposition of polysaccharides. Thus, the cell plate contains a gradient of developmental stages at any given time. A recent review by (Sinclair et al., 2022) presents an animated overview of cytokinesis.

The synchrony and precisely timed deposition of membrane material and polysaccharides control the cell plate’s expansion, stability, and maturation into a new cross wall (Drakakaki, 2015; Smertenko et al., 2017; Gu and Rasmussen, 2022). However, little is known about how these mechanisms are orchestrated (Drakakaki, 2015; Smertenko et al., 2017). Many GTPases are localized at the cell plate (Chow et al., 2008; Geldner et al., 2009; Qi et al., 2011; Qi and Zheng, 2013; Berson et al., 2014; Mayers et al., 2017; Shi et al., 2023), including the Rab-related GTPase RABA2a, which is involved in the delivery of *trans*-Golgi network derived vesicles to the leading edge of the cell plate. As such, RABA2a is considered a good marker for post-Golgi vesicles directing cargo to the cell plate (Chow et al., 2008). Necessary, vesicular fusion events during Arabidopsis cell plate formation are mediated by SNARE complexes acting redundantly in cytokinesis that involve KNOLLE and its partners (El Kasmi et al., 2013).

Our current insights into cell plate formation are based on two-dimensional, electron (Samuels et al., 1995; Segui-Simarro et al., 2004) and *in vivo* fluorescence micrographs, with only minimal contributions from multidimensional spatiotemporal data (reviewed in (Drakakaki, 2015; Smertenko et al., 2017; Sinclair et al., 2022; Geitmann, 2023). During the TN stage, callose, a β-1,3-glucan polysaccharide, is temporally integrated into the cell plate (Samuels et al., 1995). Callose is proposed to contribute as a spreading force in the stabilization and maturation of cell plate (Zawaid wt al., 2022). However, its timely delivery and contribution to the stage progression of cytokinesis remain recalcitrant. Dissecting the dynamic functions of components such as callose, other polysaccharides, and the phragmoplast machinery, is necessary in order to understand their contribution to cytokinesis. Even with the advancement in many technologies, research into cell plate development is hindered by the fact that mutations in cytokinesis such as those of callose synthase (GSL8) are often lethal (Desprez et al., 2007; Chen et al., 2009; Thiele et al., 2009) making a genetic approach to the underlying questions unfeasible. Therefore, imaging modalities that allow acquisition during the whole cytokinesis process in space and time without compromising image quality are required.

So far, confocal-based modalities are not ideal as they lead to photobleaching, making it nearly impossible to capture the entire process of cell plate from initiation to maturation. In addition, confocal-based modalities require the presence of an already initiated cell plate to focus on image acquisition with the compromise that early stages are not captured. Further, each event should be imaged separately via a tedious effort to manually correct sample drifting due to tissue growth (often at the root tip). Such studies are not easily scalable because they require significant manual input to generate datasets that allow statistical analysis. Recent technological and methodological advances are now allowing four-dimensional (4D, XYZT), *in vivo* fluorescence microscopy, which can provide more biologically relevant, quantitative information.

The development of lattice light sheet microscopy (LLSM) enables lengthy image acquisitions with significantly minimized photobleaching. LLSM is a promising avenue in dissecting mitosis across different biological systems allowing for easier acquisition of multiple cytokinesis events simultaneously, from their earliest detection, while it decreases the constant need to manually correct during acquisition (Chen et al., 2014; Aguet et al., 2016). However while this is an enabling modality, this approach has not been adopted to study cytokinesis in plants due to the efforts required to establish an imagining and quantitative analysis routine.

In this study we adopted LLSM to study cytokinesis in plants. Taking advantage of LLSM, we investigated cell plate dynamics using the cytokinesis marker RABA2a coupled with pharmacological inhibition of callose. Quantitative analysis revealed the presence of distinct phases with easily identifiable transition points during cell plate development which were altered by inhibition of callose. The presented here adopted imaging modality with quantification analysis and identified stage transition can help further untangle the complex process of cytokinesis and interrogate the biological role of its components.

## Materials and Methods

### Plant materials and growth

All Arabidopsis (*Arabidopsis thaliana*) seedlings were grown as described previously (Park et al., 2014). Briefly, seeds were sterilized for 10 min in a solution containing 10% (v/v) sodium hypochlorite (NaOCl)/ 80 % (v/v) ethanol / 10% (v/v) water followed by three washes in 90% (v/v) ethanol for 1 min and air dried. Seeds were germinated on square plates containing Murashige and Skoog (MS) medium (Sigma; one-quarter strength) supplemented with 1% (w/v) Suc, pH 5.7, solidified with 0.5% (w/v) phytagel (Sigma) and supplemented with designated chemicals. Plates were incubated at 10 ° off vertical in a growth room with a 16-h-light/8-h-dark photoperiod for 3 to 5 days. Transgenic lines expressing YFP-RABA2a (Chow et al., 2008) were used to observe subcellular phenotypes upon chemical treatment (Park et al., 2014).

### Chemical treatment and staining procedures

For microscopy, seedlings were grown as described above for 3 d and then transferred to 48-well plates containing 2 mL of liquid MS medium supplemented with 390 mM DMSO (Sigma), 50 μM endosiden 7(ES7) (ChemBridge) (or otherwise indicated concentration) and allowed to grow for two hours as pulse treatment. Chemicals were diluted to their working concentrations from 1,000× stock solutions. FM 4-64 (5 µM) diluted 1:1000 from a stock solution, applied for 5 minutes, was used to stain the plasma membrane. Aniline blue fluorochrome (Biosupplies) was used to detect callose at 0.1 mg/mL in H_2_O, diluted from 1 mg/mL stock. Staining was performed directly on imaging slides.

### Transmission electron microscopy

Four-day-old Arabidopsis seedlings, treated under 50 µM ES7 for 1 hour, were analyzed by TEM. Root tips (1mm) were excised and fixed using high-pressure freezing methods as described (McDonald, 2014; Otegui, 2020). Excised root tips were placed in a type B freezing planchette containing yeast paste as a filler/cryoprotectant and frozen under high pressure in a high-pressure freezing unit (BAL-TEC HPM 010). Freeze substitution was carried in 1% OsO4 and 0.1% uranyl acetate in acetone, followed by infiltration and embedding in Epon resin as described earlier (McDonald and Webb, 2011; McDonald, 2014). Thin sections (70 nm) were cut on a Leica Ultracut E ultramicrotome, picked up on Formvar-coated slot or 100 mesh grids, and poststained for 7 min in 2% uranyl acetate in 70% methanol, followed by 4 min in Reynold’s lead citrate. Images were taken with a Gatan Ultrascan 1000 camera on a FEI Tecnai-12 electron microscope operating at 120 kV.

### Confocal/ high resolution microscopy

#### Image acquisition: Zeiss Airyscan 980 or Leica SP8 were used for confocal imaging

Leica SP8 was used to image YFP-RABA2a seedlings co-stained with FM 4-64 (5 µM) under a 100X/1.4 NA oil objective HC PL APO CS2 employing high resonant scanning with line averaging set to 15. Excitation at 512 nm was used for both YFP-RABA2a and FM 4-64, with emissions collected at 520-558 nm and 650-800 nm for YFP and FM 4-64, respectively. For the time-lapse series, Z-stacks were collected at intervals of ∼1 min for 30 – 45 min. Images are representatives of 5 biological replicates (independent seedlings).

Zeiss 980 was used to image YFP-RABA2a seedlings co-stained with Aniline Blue at single time points, using Airyscan Super Resolution (SR)8Y mode. The fluorescent signal of YFP-RABA2a was exited using a 514nm laser at 25% power and aniline blue fluorochrome using 405nm laser at 15% power. All images were collected using the LD LCI Plan -Apochromat 40X/1.2 NA (Korr DIC M27) water objective. Images are representatives of 5 biological replicates (independent seedlings).

#### Image processing

Leica SP8 data were deconvolved using Classic Maximum Likelihood Estimation. (CMLE), manually adjusting for background with a maximum of 20 iterations and corrected for XYZ drift using Huygens (SVI). Zeiss 980 Airyscan collected images were processed using the Zeiss built-in deconvolution software using standard settings. All data were exported to Imaris, and Bitplane for segmentation and 3D or 4D visualization. Figures were assembled using Affinity Designer.

### Lattice Light Sheet Microscopy

#### Imaging

Imaging was performed using the LLSM at the Advanced Imaging Center (AIC), Janelia research campus, following sample preparation and imaging procedures as previously described (Chen et al., 2014). Arabidopsis seedlings 3 to 4 days old were mounted on a 5 mm coverslip Warner Instruments Cat. No 64-0700 (CS-5R). One seedling was mounted per coverslip within a thin layer of 0.5% low melting agarose (w/v) in H_2_O. The cover slip was clipped to the end of a long extension of the sample holder and submerged in a bath filled with 8.5 mL ¼ MS liquid media. The opposite end of the coverslip holder was bolted to the sample piezo (Chen et al., 2014). YFP-RABA2a plants were stained with FM 4-64, and both fluorophores were excited using a 488 MPB fiber laser. A two-camera system, 2x Hamamatsu Orca Flash 4.0 v2 sCMOS, was used to acquire the images of the two fluorophores.

#### LLSM Image processing

All data were acquired in the dithered mode and were deconvolved by using a Richardson-Lucy algorithm adapted to run on a graphics processing unit (GPU) (NVIDIA, GeForce GTX TITAN), using an experimentally measured point spread function (PSF) for each emission wavelength. Before visualization, all 3D data sets acquired via sample scan in the x, y, and z coordinated system were transformed (“deskewed”) to the more conventional x, y, and z coordinates in the GPU.

### Selection of points of interest for channel alignment and image stabilization

#### Channel alignment

The BigStitcher plugin in ImageJ was used to process data sets before quantitative analysis. All files using BigStitcher were converted to .xml to register how the data is stored and track any internal processing following the pipeline described in advanced imaging center (AIC) guidelines: https://knowledge.aicjanelia.org/posts/20200730-stabilize-roi-selection/. One .xml file was generated per time-lapse data set. For our experiments, the metadata denoted FM 4-64 as channel 0 and YFP-RABA2a as channel 1.

Once a file was generated, channel alignment was applied first, followed by drift correction (movement in the field of view). Note that the drift correction can lead to misalignment if alignments are not applied in this sequence. Channel alignment followed a similar routine to the drift correction as described in AIC guidelines: https://knowledge.aicjanelia.org/posts/20200730-stabilize-roi-selection/ with the following modifications listed below.

1. In the Multiview Explorer window, the first time point was selected for channel 0 (FM 4-64), and the command ‘Detect Interest Points’ was navigated for the type of interest points “Difference-of-Gaussian” with no other restrictions selected. This was repeated for channel 1 (YFP-RABA2a).
2. Then the parameters used to identify the interest points were refined using the sigma and threshold sliders. The minimum threshold was used to detect the space within each cell enabling individual selections of interest for each cell. Sigma values were set to a range of 8-9 to obtain detectable *points/regions* inside each cell that can be used for the alignment. Adjustments were made to create the best recognizable points common between the two channels to create the best alignment in step 3.
3. The points of interest were registered using a precise descriptor-based (translation invariant) algorithm to align the two channels and validate through the viewer window 3D function. The transformation model is rigid with an allowed error for random sample consensus iterative methodology known as RANSAC (px) of 7-8. This was repeated with several alterations to the sigma threshold until the two channels overlapped.
4. Once complete, the BigStitcher windows were closed, and the Plugins menu was navigated to the following sequence: Multiview Reconstruction -> Batch Processing -> Tools -> Duplicate Transformations. Transform one-time points to other time points.

The transformation was applied to other time points across the data set.

#### Drift correction

Following channel alignment, drift correction was resolved using the steps below as described in AIC guidelines: https://knowledge.aicjanelia.org/posts/20200730-stabilize-roi-selection/. Note the significant differences from the channel alignment steps: 1) the use of all time points in one channel, instead of the first time point and 2) the adjustment of Gaussian values to identify the cell wall corners across the time points, instead of the space inside the cells that was used for channel alignment. The threshold was set to the maximum, while the sigma value was set to a range from 7.5-9.5.

#### Create a bounding box for region of interest

Region of Interest (ROI) selection and export as 3D .tiffs were used to decrease file size and obtain workable files by following the steps “Create Bounding Box for Region of Interest” as outlined in AIC guidelines: https://knowledge.aicjanelia.org/posts/20200730-stabilize-roi-selection/.

#### Image stack assembly and bleach correction

The .tiff files generated from the Bounding Box ROI step were then separated into folders by each channel and concatenated into full-time series using ImageJ. The time series were then bleach-corrected using histogram matching. Finally, processed data were first exported to individual .tiffs and subsequently to .ims using the Imaris file converter. Before converting, the settings were adjusted for the voxels, XYZT frame, and image acquisition time points. Files were then finalized and imported into IMARIS Bitpane for 4D volumetric rendering. All time series at this step contained two data sets, one “raw” data and one for the histogram-matched bleach-corrected.

### Imaris Segmentation

Surface rendering of individual cell plates was performed using the surface creation tool using Imaris x 64 9.6.0 and the steps of filament tracer https://qbi.uq.edu.au/research/facilities/microscopy-facility/image-analysis-user-guides/analysis-software/imaris/creating-surface-imaris … with modifications as outlined below:

1. In the first step of the surface creation the settings were left open, with all options unselected, unless a specific ROI was first separated. Using the ROI tool, a bounding box enabled the separation of a specific region and further reduced the working file size.
2. Surface segmentation was performed using the “new surfaces” function. The appropriate channel (histogram corrected) was used. Surface detail was set to 0.1 µm, and thresholding was based on background subtraction with the values set based on the diameter of the largest sphere fitting the object. To obtain these values, the diameter of the largest shape was measured in the 2D view under the slice mode.
3. During the next step, a histogram of image intensity was adjusted to best fit the objects of interest. Multiple surfaces/segmentations could be used for different cell plates and different time points for statistical analysis.
4. In the next step, a preview was generated by Imaris with final surfaces over all the objects that were detected in the ROI. Adjustments were made on the number of voxels to better refine the observed objects. Voxel set to 20 throughout the data set allowed for the best observation of our data. The “autogestion motion” function was implemented with the drift correction above a max distance of 4 µm and max gap size of 3 µm.
5. Following surface rendering, individual cell plates were segmented by selecting their corresponding sub-surfaces and rebuilding the cell plate individual tracks and connecting the sub-tracks together. Surfaces and volumes of cell plates were extracted, uniquely named, and quantitatively analyzed. We note that working with different channels separately leads to manageable file sizes within the processing computer power. As the different channels are aligned, they can be later combined after further processing.
6. Cell plate diameter values were recorded manually for each time point during cell plate formation by measuring the largest distance of YFP-RABA2a. The FM4-64 plasma membrane stain was used to determine the predicted final cross-wall width and normalized the diameters, accounting for cell size variations (Chow et al., 2008; van Oostende-Triplet et al., 2017). Finally, normalized diameter values were averaged at their respective time points across cell plates.

### Quantification and statistical analysis

The data generated from the intensity based IMARIS algorithms were compiled within Excel files and further analyzed. The Origin2021 software (Origin. 2003. Origin 7.5. OriginLab Corp., Northampton, MA) was used to normalize the data to their maximum volume or area. To determine an approximate rate of change of measured volumes/areas, the data was fitted to a polynomial (Adjusted R-Square 0.69 ± 0.03), on which derivatives were drawn. For visualization purposes, the derivatives were linearly shifted, creating a time when all rates passed through zero. However, raw datasets were considered for fitness comparison and any further statistical analysis differences.

### Binning approach for further statistical analysis

For **Fig.S1**, the data were divided into five groups (bins) for further statistical analysis. These bins correspond to the normalized volume changes based on the following approach: bin 1 (increasing values from 0 - 0.33 µm^3^), bin 2 (increasing values from 0.33 - 0.66 µm^3^, bin 3 (increasing values from 0.66 −1 µm^3^), bin 4 (decreasing values from 1 - 0.66 µm^3^), bin 5 (decreasing values from 0.66 - 0.33 µm^3^), with bin 6 (decreasing values from 0.33 µm^3^ until reaching the final minimal RABA2a cell plate volume. The width of the bins, as well as the divide between increasing and decreasing values, were chosen for simplicity. The derived rates from each of the volume bins were then averaged. These averages were statistically analyzed using two-tailed pairwise t-tests (GraphPad) and then were plotted as shown in the **Fig. S1**.

P-values for diameter studies were additionally calculated using a second binning approach. The normalized diameter values were averaged across cell plates at their respective time points (**Fig. 6A**). For comparisons, The ES7 time points were laterally adjusted such that the averaged ES7 values at the starting point matched the corresponding control value (**Fig. 6C**). P-values were then calculated using pairwise t-tests (GraphPad) (**Fig. 6D**).

## Results

### Development of a 4D Lattice Light Sheet Microscopy image acquisition and processing pipeline to study plant cytokinesis

Cell plate assembly, expansion, and maturation are regulated in space and time over an extended period. Therefore, 4D imaging is necessary to examine in depth the membrane and cell plate expansion dynamics. We first used confocal microscopy equipped with a resonance scanner, allowing for quicker acquisition and a short exposure time to the excitation laser, to minimize photobleaching over a long period of recording cytokinesis. Using YFP-RABA2a (Chow et al., 2008) as a cytokinesis marker, imaging cell plate development in high-resolution was tractable (**Video. S V1,**). Time lapse imaging showed centrifugal expansion of cell plate. This approach, however, was not sustainable for the dissection of a large number of cytokinesis events. Confocal microscopy could only manually track developing cell plates, one at a time, limiting scalability. In addition, using this approach, only previously sufficiently assembled cell plates could be imaged since their presence was necessary to be identified in the first place.

To circumvent these hurdles and capture full cell plate events while improving throughput, we used LLSM. The LLSM affords imaging at much faster rates and with less light exposure of the sample, which minimizes photobleaching and phototoxic effects (Chen et al., 2014). LLSM helps extend the observation time while affording higher axial resolution with similar lateral resolutions to those obtained with confocal microscopy. Using the LLSM, we recorded the marker YFP-RABA2a in the root tips of 3-day-old *Arabidopsis thaliana* seedlings. We generated 4D data capturing the complete process of cell plate development with minimal photobleaching, allowing the acquisition of data sets amenable to in-depth quantitative analysis. The cell membrane and cellular architecture dye, FM 4-64, was used to provide a reference for overall cell shape and cell plate expansion perspective. Once image acquisition was completed, a pipeline was developed primarily using open-source software for image processing followed by quantitative analysis (**Fig. 2**, more details in the material and methods section). In brief, this involved image deskewing, aligning the two fluorescence channels, corrections for (XYZ) drift, and selection of a ROI to obtain manageable data size. Once data was exported to readable formats (.tiff files), they were concatenated into time series stacks for each channel. The image files were bleach-corrected by histogram matching across time points for the whole stack (**Fig. 2B &C**). Segmentation of developing cell plates was performed using an intensity value-based algorithm in IMARIS, followed by quantitative analysis of cell plate volume, surface area, and diameter (**Fig. 2D**).

**Fig. 1.**
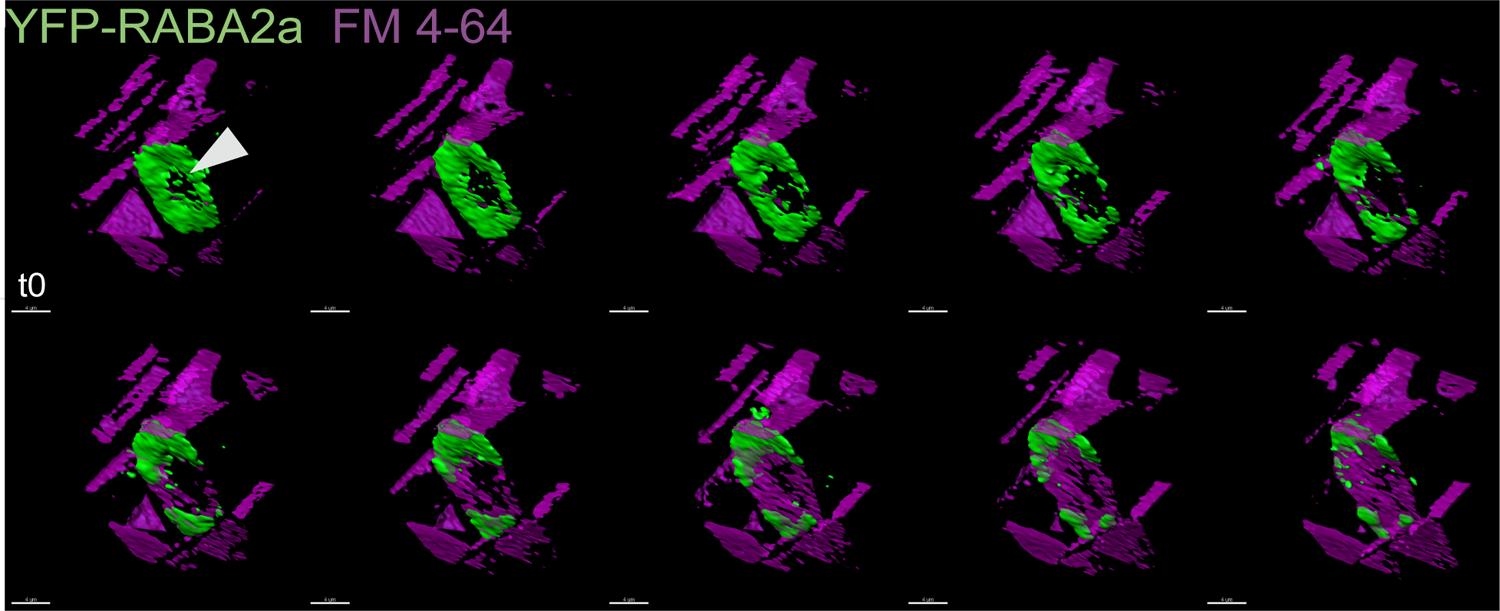
YFP-RABA2a dynamics at the cell plate. YFP-RABA2a (green) vesicle accumulation and FM 4-64 (purple) stained plasma membrane show the transition from vesicle accumulation to mature membrane throughout cell plate development in untreated plants. The accumulation of YFP-RABA2a at the cell plate periphery during the maturation with a concurrent increase in the membrane content in the center shows centrifugal growth and maturation. Arrow indicates RABA2a at the cell plate. Data was collected on a Leica SP8 STED microscope. Scale bar = 10 µm. Δt = 1 min.

**Fig. 2.**
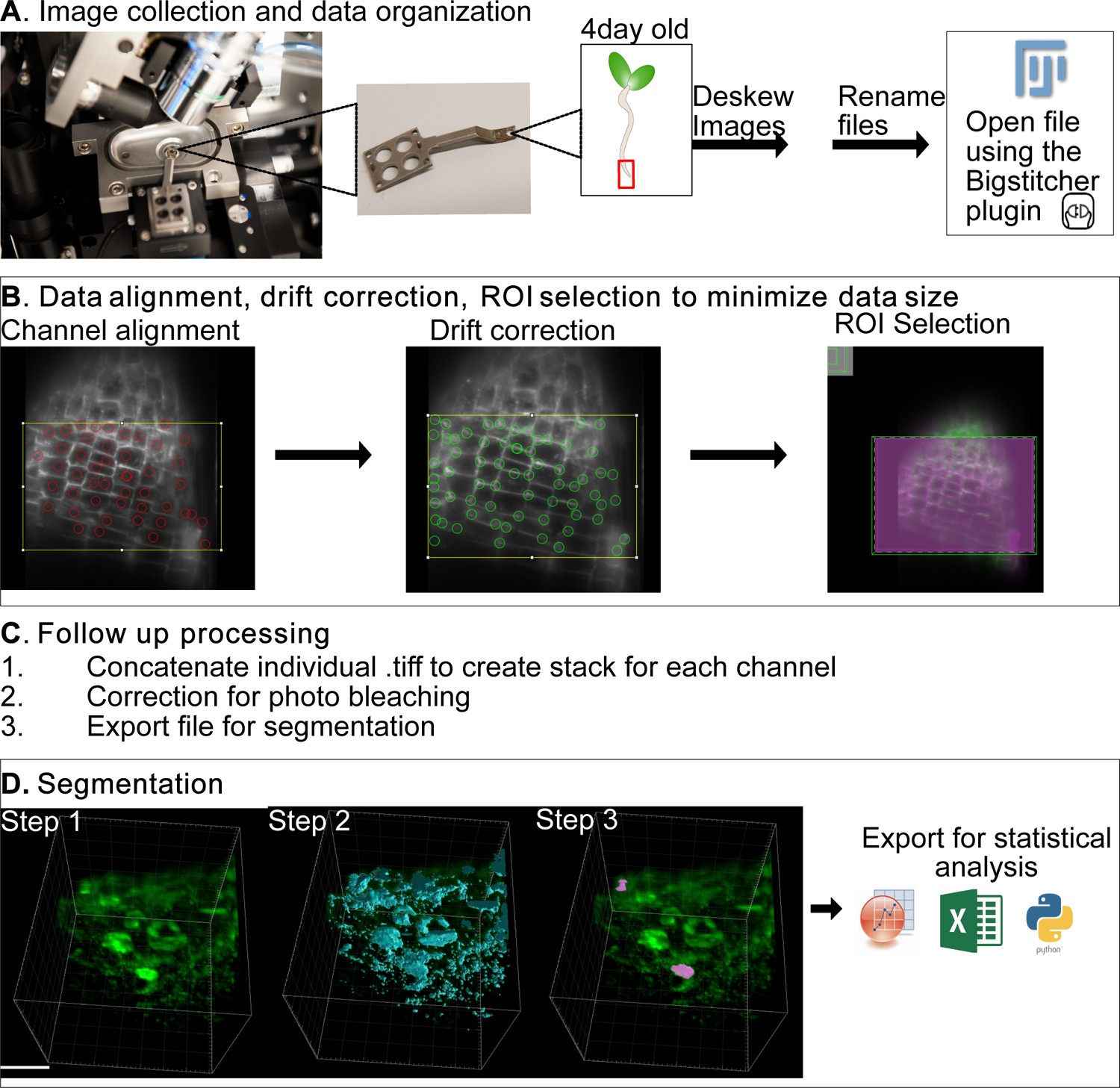
Schematic representation of Lattice Light Sheet data processing pipeline. Schematic representation of the processing workflow for collecting plant cytokinesis events under Lattice Light Sheet Microscopy (LLSM). The workflow is separated into four overarching steps: **(A)** image collection and data organization, **(B)** data alignment with drift correction and ROI selection, **(C)** concatenation of each time point to create a Z stack and bleach correction follow-up processing, **(D)** image collection and data organization. Scale bar = 20 µm.

### Quantification of cell plate dynamics shows three district developmental stages based on membrane accumulation patterns

With the adoption of LLSM, cell plate growth from its point of origin at the vesicle accumulation stage until completion of cytokinesis can be observed, a time frame that was not previously established due to constrains in data sets via confocal and electron microscopy (Segui-Simarro et al., 2004; Higaki et al., 2008; van Oostende-Triplet et al., 2017; Geitmann, 2023). An example is presented in **Fig. 3A, a** pink segmented surface, in which volume and area change and rates are tracked by a bolded blue line in **Fig.s 3B-E**). As shown **in Fig. 3A, B**, completion of cytokinesis takes place within 20-30 minutes. Within this time window, most cell plate volume peaked during the first 8 minutes (**Fig. 3A**). Then the cell plate expanded laterally and flattened with the gradual reduction of the vesicle accumulation marker dispersing at the cell plate rim and finally disappearing (**Video. S V1**), likely representing transitions to discontinuous phragmoplast stages (Smertenko et al., 2017; Sinclair et al., 2022). With LLSM, observation of multiple cytokinetic events within the same root tip was possible. However, there was no apparent trend associated with each root tip. Thus, each cell plate was treated as a separate event.

**Fig. 3.**
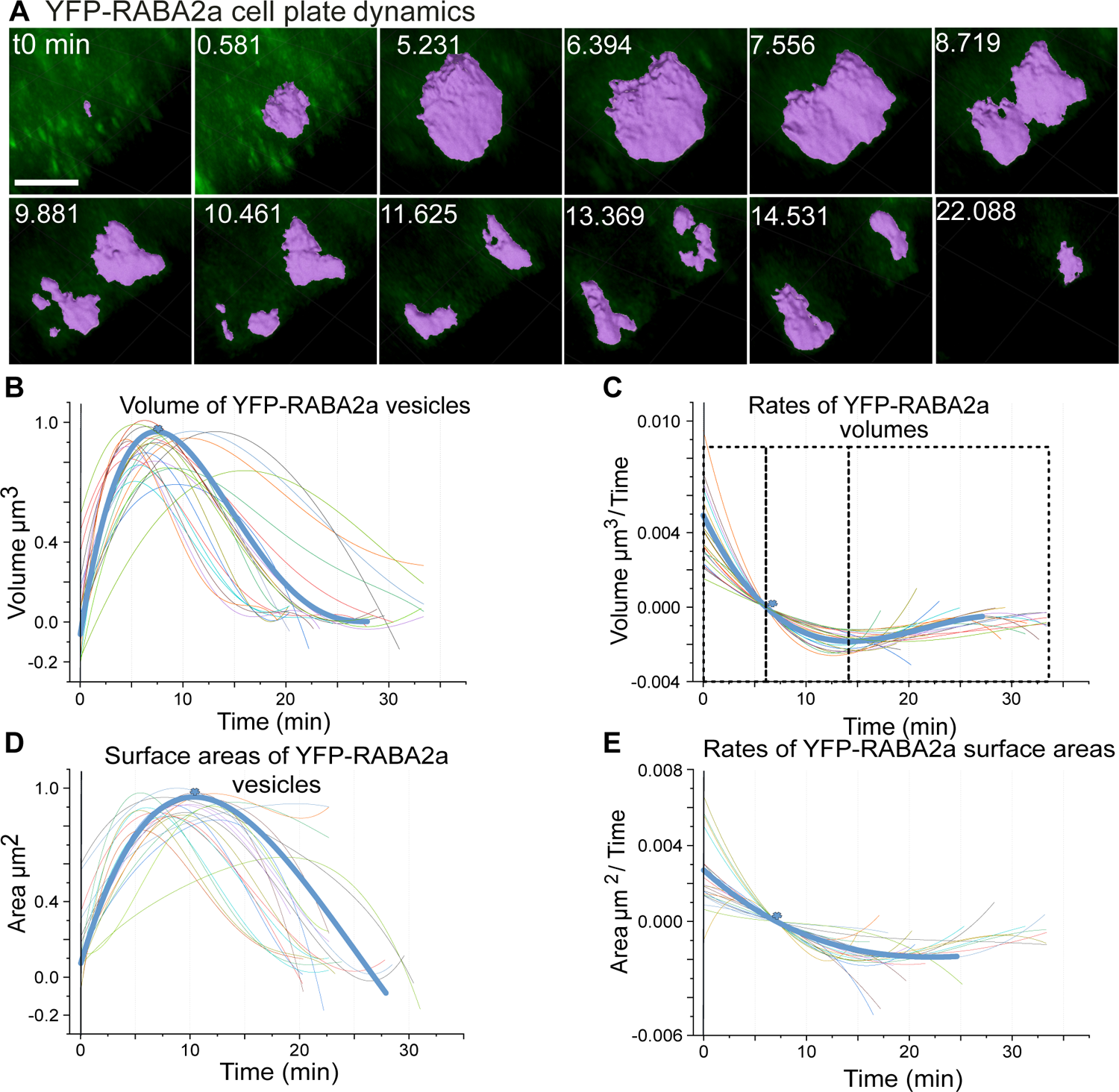
Quantitative YFP-RABA2a dynamics at the cell plate. **A)** Representative snapshots of a time series showing the transition of YFP-RABA2a (green) in segmented cell plates (purple) during their expansion and maturation. **B-C)** Volumes of segmented cell plates and their growth rates. **D-E)** Surface areas of segmented cell plates and their rates of change. Each line indicates an individually segmented cell plate. Note the cell plate shown in **(A)** is bolded in all graphs. Normalized volumes and areas, respectively, of developing cell plates were fitted to a second-degree polynomial distribution **(B&D)**. Additionally, the rates of change were derived using first-degree derivatives of the polynomial fits for both the volume and the bounding surface, respectively. Scale bar = 5 µm.

A quantitative analysis of segmented cell plates based on the cytokinetic marker was performed to understand better the spatiotemporal dynamics of vesicle accumulation and cell plate expansion. Normalized volumes of developing cell plates were fitted to a second-degree polynomial regression, allowing accumulation trends to be observed across biological samples (**Fig. 3B**). As aforementioned, the cell plate volume accumulation based on RABA2a peaked within ∼8 minutes, followed by a rapid reduction over the following 8 minutes. The rates of volume (1.54×10^3^-9.45 x10^3^ µm^3^/min) accumulation determined by the first-order derivative showed a net positive addition of RABA2a vesicle/membrane material for the first 8 minutes (**Fig. 3C**). Within this period, a short interval corresponds to linear growth. The first 8 minutes marked a turning point in which the rates of volume accumulation switched to a net negative value **(Fig. 3C, asterisk**). The negative rate values likely represent a large amount of recycled material during cell plate expansion and maturation, as predicted by electron microscopy analysis (Otegui et al., 2001; Segui-Simarro et al., 2004). We also quantified the bounding surface area corresponding to the segmented cell plates. The accumulation of RABA2a surface area and their corresponding rates (**Fig. 3D, E**) showed an overall similar trend with volume accumulation.

In summary, three distinct phases became apparent based on our quantification: phase I, a rapid phase of cytokinetic vesicle material delivery with a positive RABA2a cell plate volume rate, followed by a substantial volume reduction (phase II). Finally, a third phase (phase III) of minimal cytokinetic vesicle presence at the rim of the cell plate, took place before joining the parental cell wall.

### Inhibition of callose deposition arrests cytokinesis and induces aberrant membrane accumulation patterns

Our data show that while vesicle volume accumulation is a very rapid process, the expansion and maturation phase is lengthier. The latter stages are accompanied by a significant loss of cell plate volume, likely due to the recycling of membrane material. During the transition of the cell plate from a membrane network to a fenestrated sheet, the accumulation of the polysaccharide callose is implicated in stabilizing this network and providing a spreading force for expansion and maturation (Samuels et al., 1995; Jawaid et al., 2022). To better understand the role of callose during these dynamic cell plate transitions, we applied Endosidin 7 (ES7), a chemical that explicitly inhibits cytokinetic callose (Park et al., 2014, Drakakaki et al., 2011).

Observed by confocal microscopy, RABA2a accumulation was not affected by the 2-hour ES7 pulse treatment. However, the cell plate expansion and maturation were impaired, leading to stagnation, shown by the cytokinesis marker YFP-RABA2a and FM 4-64 staining (**Fig. 4A**). We then used transmission electron microscopy (TEM) on high-pressure fixed root tips to resolve changes induced by ES7 at the ultrastructural level. As shown in **Fig. 4Bi**, the structure of cytokinetic vesicles was not affected during the accumulation stage, and no discernable aberrations in the vesicle or Golgi morphology were observed. However, irregular cell plate stubs were observed in later stages of cytokinesis (**Fig. 4Bii, arrows**) in contrast to normally expanding cell plates as previously reported in Arabidopsis (Samuels et al., 1995), verifying our light microscopy confocal observations. While informative, confocal and electron microscopy were not ideal for studying the dynamic nature of cell plate development under ES7 treatment.

**Fig. 4.**
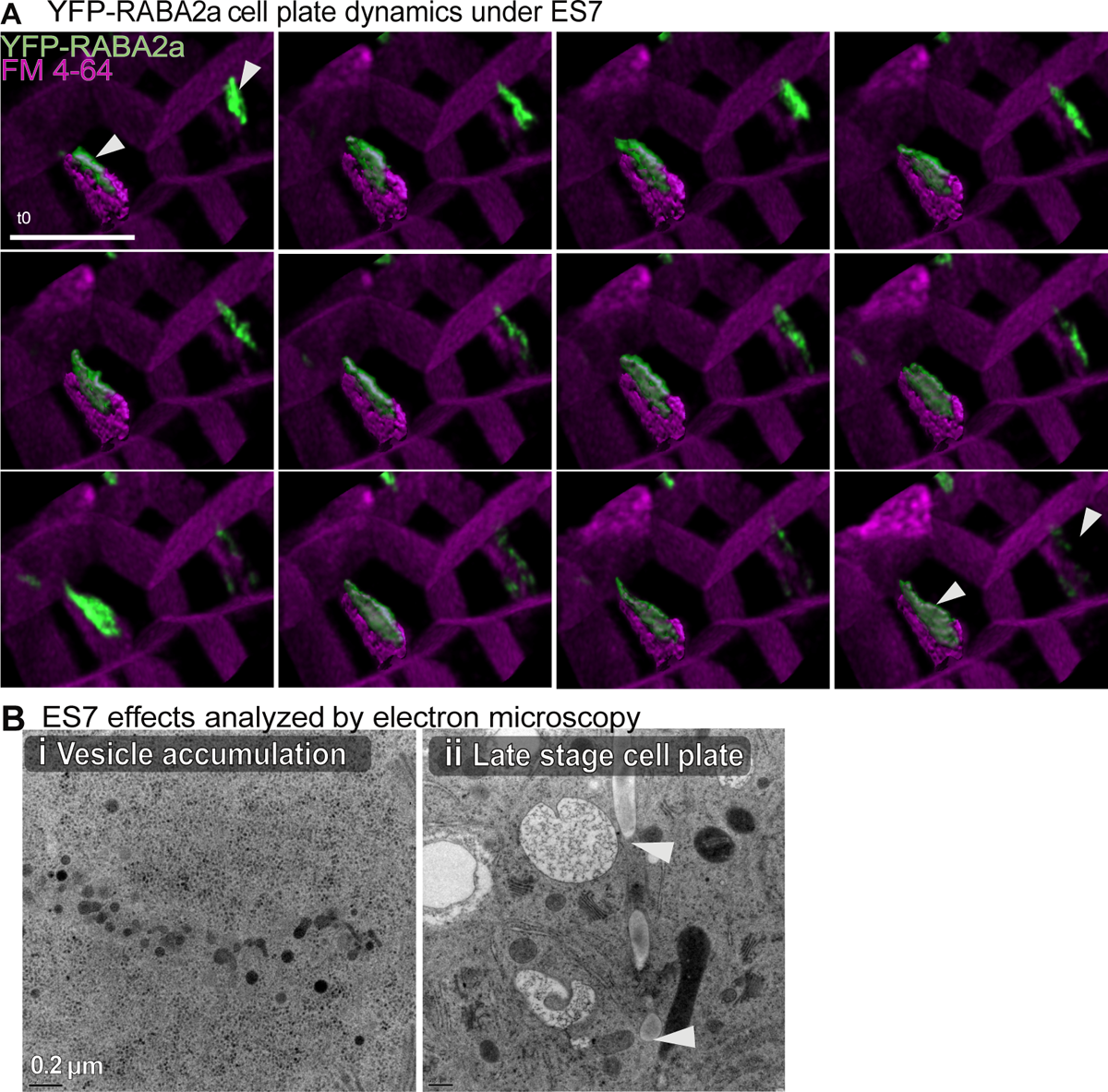
YFP-RABA2a cell plate dynamics under Endosidin 7 treatment. **A)** YFP-RABA2a (green) vesicle accumulation and FM4-64 (Purple) stained plasma membrane show cell plate development under Endosidin 7 (ES7) treatment. Note the abnormal pattern and the fragmentation of the cell plate as it transitions during different time points. The pattern of YFP-RABA2a at the cell plate periphery does not expand radially, as seen in the untreated plants. Further, the cell plate cannot follow normal maturation into membranes detectable with FM4-64. Data was collected on a Leica SP8 STED microscope. Scale bar = 10 µm. Δt = 1.5 min. **B)** Observed ES7 effects using electron microscopy. No effect was seen in the vesicle accumulation stage **(I)**, but cell plate fragmentations were observed in the late stages **(II)**. Arrows indicate late stage cell plates affected by ES7. Scale bar in Bi = 0.2 µm. Scale bar in Bii = 0.5 µm.

### Membrane transition between cytokinesis stages is affected by the loss of callose

Again, highlighting the confocal and electron microscopy offer valuable information, they have some limitations when studying the dynamic nature of cell plate development under ES7 treatment. Thus, we again employed LLSM. The application of LLSM still allowed imaging of various cytokinetic events, facilitating quantification and providing an analysis of the impact of callose inhibition. It’s worth noting that the observed events took longer compared to untreated plants. Overall, LLSM proved to be a valuable tool for capturing essential data on cell plate development under ES7 treatment providing an overview of the effect of callose inhibition.

In contrast to the untreated ones (**Fig. 3**), the cell plates subjected to a 2-hour ES7 pulse treatment (**Fig. 5**) exhibited a notable accumulation of RABA2a volume that persisted beyond 15 minutes without significant expansion in the cell plate itself. Meanwhile, the marker maintained a condensed structure, eventually fragmenting throughout the lifetime of imaged cell plates (**Fig. 5A & Video. S2),** cell plates indicated by blue). Another ES7-induced phenotype, though less frequent, showed relative expansion in the cell plate but also eventually fragmented in an erratic pattern (indicated by pink **Video. S3)**, cell plate indicated by pink and blue for both phenotypes). Overall, the relative expansion did not match that observed in untreated samples as further discussed below. Under the influence of ES7, the attempted cell plate development ran longer than controls, averaging 35 minutes ±13.7 min compared to the 20-30 minutes in untreated samples. The variation in ES7-induced phenotypes was likely due to the timing of the drug’s effect at different cytokinetic stages, as multicellular root tips are not synchronized.

**Fig. 5.**
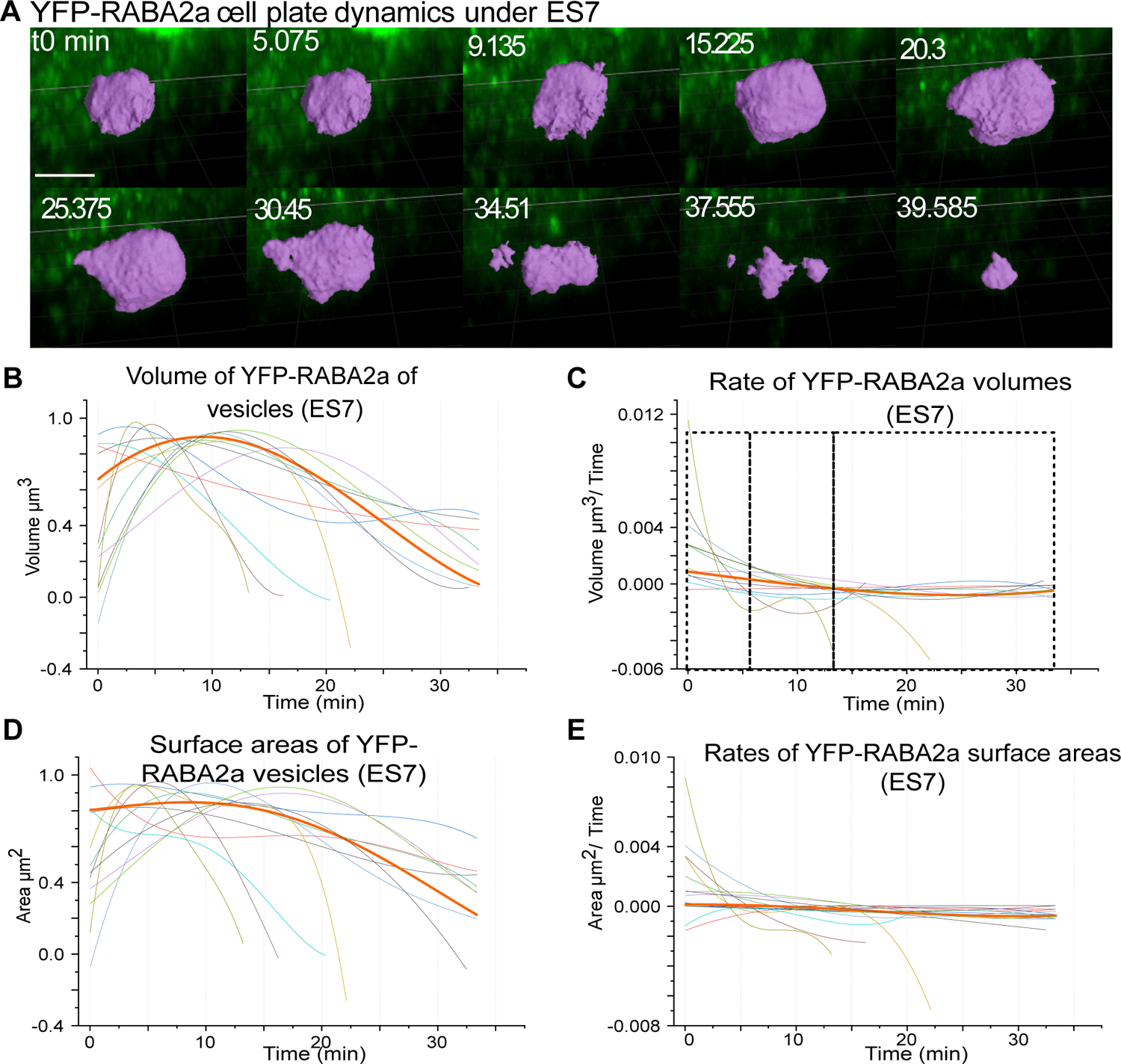
Quantitative YFP-RABA2a dynamics at the cell plate under ES7 treatment. **A)**. Representative snapshots of a time series showing the YFP-RABA2a segmented cell plate transition under ES7 treatment. The segmented cell plate is shown in purple from its first emergence, not able to expand radially and flatten. **(B-E)** Volumes and bounding surfaces of segmented cell plates and their rates of change. Each line indicates an individually segmented cell plate. Note that the cell plate shown in **(A)** is bolded in all graphs. Normalized volumes and bounding surface areas of developing cell plates were fitted to a second-degree polynomial distribution **(B&D)**. First-degree derivatives present the corresponding rates of change of the polynomial fits for the volume and the bounding surface area, respectively (C&E). Scale bar = 10 µm.

Quantifying RABA2a cell plate volumes (**Fig. 5B**) and their corresponding rates of change (**Fig. 5C**) under a 2-hour ES7 pulse treatment showed an initial phase of rapid volume growth, followed by a slower to stagnant accumulation of RABA2a membrane material, leading to a prolonged plateau of retained volume over a longer timeframe. This prolonged plateau contributes to the increased average division time compared to the untreated plants (**Fig. 5B, C**). Similarly, the exterior surface area of the cell plates exhibited very similar dynamics and behavior, with a broad plateau of surface area over time and a severely depressed rate of surface area change (**Fig. 5D, E**). Overall, the phenotypic observations and quantifications of cell plate characteristics demonstrate a stark difference between ES7-treated and control cell plates. While the control cell plates followed distinct, consistent growth phases based on volume changes, marked by a transition from positive to negative rates, no apparent pattern changes in ES7-treated volumes, corresponding to discernable phase transitions, were observed.

In order to assess the impact of ES7, the volume growth rates of both control and ES7-treated cell plates were divided into equidistant volume intervals using 20% intervals (6 “bins”) for statistical analysis. We focused on five bins, as the sixth bin at the end of the collection showed high variability. The first three intervals (bins 1-3) correspond to the normalized increasing volumes in the control, ranging from 0 - 0.33 μm^3^, 0.33 - 0.66 μm^3^, and 0.66 - 1 μm^3^, respectively. Intervals 4 and 5 represent reducing volumes ranging between 1 - 0.66 μm^3^ and 0.66 – 0.33 μm^3^ respectively (**Fig. S1F).** Bin 1 (**Fig. S1A**) corresponds to the interval with maximum volume accumulation, which is similar between the control and ES7 (**Fig. 3 and 5**). Intervals 2, 3 and 5 exhibited statistically significant differences between the control and ES7 treatments (**Fig. S1B-C, E**). The most pronounced difference was observed in interval 5 (**Fig. S1E**), in which ES7-treated cell plates showed minimal growth reduction and maintained a positive average rate, in contrast to the non-treated ones. Notably, the maximum amount of volume or surface area accumulated by RABA2a throughout all the events was not statistically different between ES7 and controls (**Fig. S2**), indicating that vesicle accumulation remains unaffected by the inhibition of callose. Taken together, the data suggest that inhibition of cytokinetic callose disrupts the phase transition from cytokinetic vesicle/volume accumulation to volume reduction and cell plate maturation.

### Expansion rate analysis identifies a critical point for callose deposition

Prior models have relied on radial expansion to stage cell plate maturation, as obtained from confocal microscopy data (Higaki et al., 2008; van Oostende-Triplet et al., 2017). To gain deeper insights into the morphological stages of cell plate development and how they are impacted by callose inhibition, we also adopted the cell plate diameter as a measure of expansion (**Fig. 6, Fig. S3**). At each time point, we measured the diameter at the longest distance across the outermost opposing edges of detectable RABA2a at the cell plate’s rim, using the FM 4-64 plasma membrane stain to determine the predicted final cross-wall width and normalized the diameters, accounting for cell size variations (Chow et al., 2008; van Oostende-Triplet et al., 2017). The diameter analysis (**Fig. 6A**, **Fig. 3A, C**) displayed a logarithmic expansion pattern in control plants. In contrast, the ES7-treated plants exhibited unstable growth with a reduction trend observed after 10-12 minutes (**Fig. 6A, Fig. S3B, D**). These findings provide valuable insights into the effect of callose inhibition on cell plate development and its impact on the expansion dynamics of the cell plates.

**Fig. 6.**
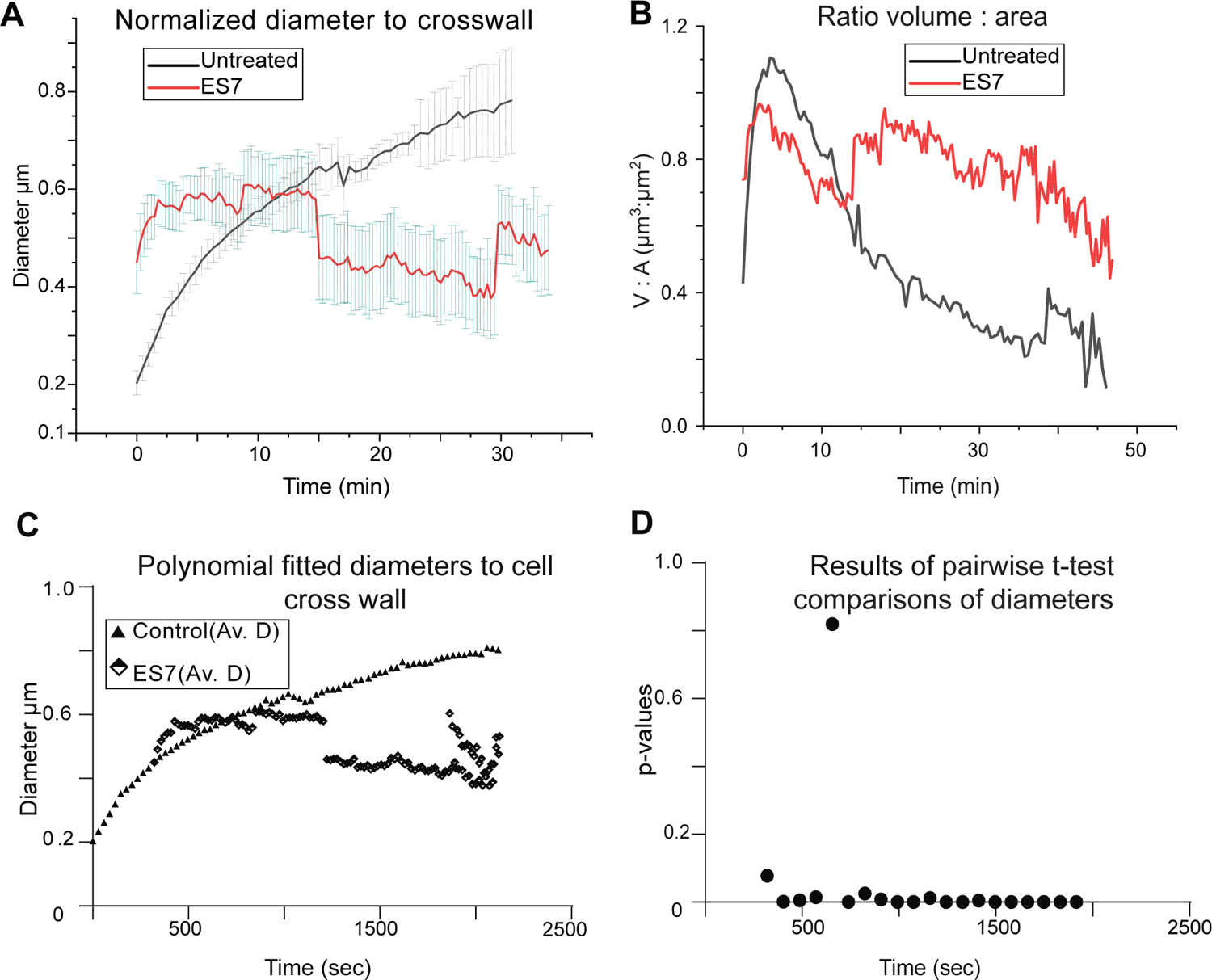
Cell plate diameter expansion under control and Endosidin 7 treatment. Cell plate transition based on cell plate diameter expansion under control and ES7 treatment. **A)** Control cell plates show a logarithmic expansion pattern, while ES7 treatment causes a more level pattern. **B)** Cell plate diameters are normalized to the expected cross-wall length. Note the intersection between control and ES7 treatment, indicating a critical point for cell plate maturation. **C)** First-degree polynomial fit was applied to cell plate diameters that have been normalized to the corresponding cross wall. **D)** Results of pairwise t-test comparisons along moving averages of **C)**. Analysis shows all but one time points are statistically different on cell plate expansion when comparing control with the absence of callose via ES7 treatment.

Interestingly, during the first 8-10 minutes that marked the bulk of vesicle volume accumulation, both ES7 treated, and control cell plates reached the same expansion levels corresponding to ∼ 60% of the predicted final diameter. While control cell plates continued growing exponentially, ES7-treated cell plates showed minimal expansion beyond the first 10 minutes (**Fig. 6A**). Based on these data, the intersection of the two growth curves indicates a critical point at which callose is essential for cell plate expansion. Notably, this time interval corresponds to the cell plate phase transition from phase I to phase II, during which volume growth changes from a positive to a negative rate and a significant volume loss occurs (**Fig. 3B, C**).

Our previously developed biophysical model highlights the importance of a spreading force, likely by the deposition of callose, for a cell plate to transition from a vesicular membrane network to a fenestrated sheet and finally to a mature cell plate (Jawaid et al., 2022). Combined with the presented data, we hypothesize that cell plate formation is characterized by a distinct, callose-dependent transition that changes from membrane accumulation to membrane recycling within approximately a 10-minute mark. We further suggest that this callose-dependent transition is due to its proposed function in providing a spreading force for cell plate expansion, in agreement with our biophysical model (Jawaid et al., 2022). Suppose that some ES7-treated cell plates achieve a level of expansion, probably due to the timing of treatment. However, the majority tend to collapse or fragment and fail to reach the parental cell wall.

There were significant technical difficulties in reliably measuring the thickness of the somewhat irregular cell plates, which led us to use instead the ratio of the cell plate volume divided by the bounding cell plate surface to estimate plate thickness. The volume-to-surface area ratio in control cell plates shows a bell curve distribution (**Fig. 6B**), with an initial volume accumulation phase followed by flattening, thinning, and diametric expansion of the cell plate. However, when treated with callose-inhibiting ES7, the volume-to-surface ratio followed a fluctuating flat line without a significant volume loss (**Fig. 6B**). This behavior might be indicative of a mechanism that critically inhibits “radial” expansion and potential redistribution/recycling of membrane material. Polynomial fitted cell plate diameters were grouped into 20 intervals (“bins”) which were subjected to statistical analysis comparing control versus ES7 treated cell plates (**Fig. 6C**). All but one interval was statistically different (**Fig. 6D**). Notably, this one interval identifies the intersection of the ES7 and control cell plate expansion curves and a critical point in which callose’s presence is essential.

### Callose deposition appears in the late cytokinetic stages

Next, we examined the localization of callose in relation to RABA2a to test if its presence corresponds to the predicted critical point during cell plate expansion. Using the callose-specific stain aniline blue fluorochrome (Evans and Hoyne, 1982) and employing multichannel live cell imaging, we characterized the transient presence of callose in relation to the cytokinesis marker RABA2a. During the early stages of cell plate development, no callose signal was identified, **Fig. 7A**, while significant RABA2a accumulation was observed, marking an expanding cell plate. During later stages of cell plate development, callose was clearly identifiable and spatially distinct from RABA2a. The cytokinetic vesicle marker RABA2a showed a ring or partial ring pattern (**Fig. 7C, D**), while callose was observed in an expanded disk form (**Fig. 7C-E**). We observed callose deposition beginning at this critical morphological transition point corresponding to the ∼8-10-minute mark characterized by a change from a disk-shaped to a ring-shaped RABA2a pattern. We conclude that the polysaccharide deposits during the transitional stages of RABA2a volume reduction and cell plate expansion. This pattern distinctly corresponds to phases II and III and covers the interval beyond the crossover point predicted by cell plate diameter expansion.

**Fig. 7.**
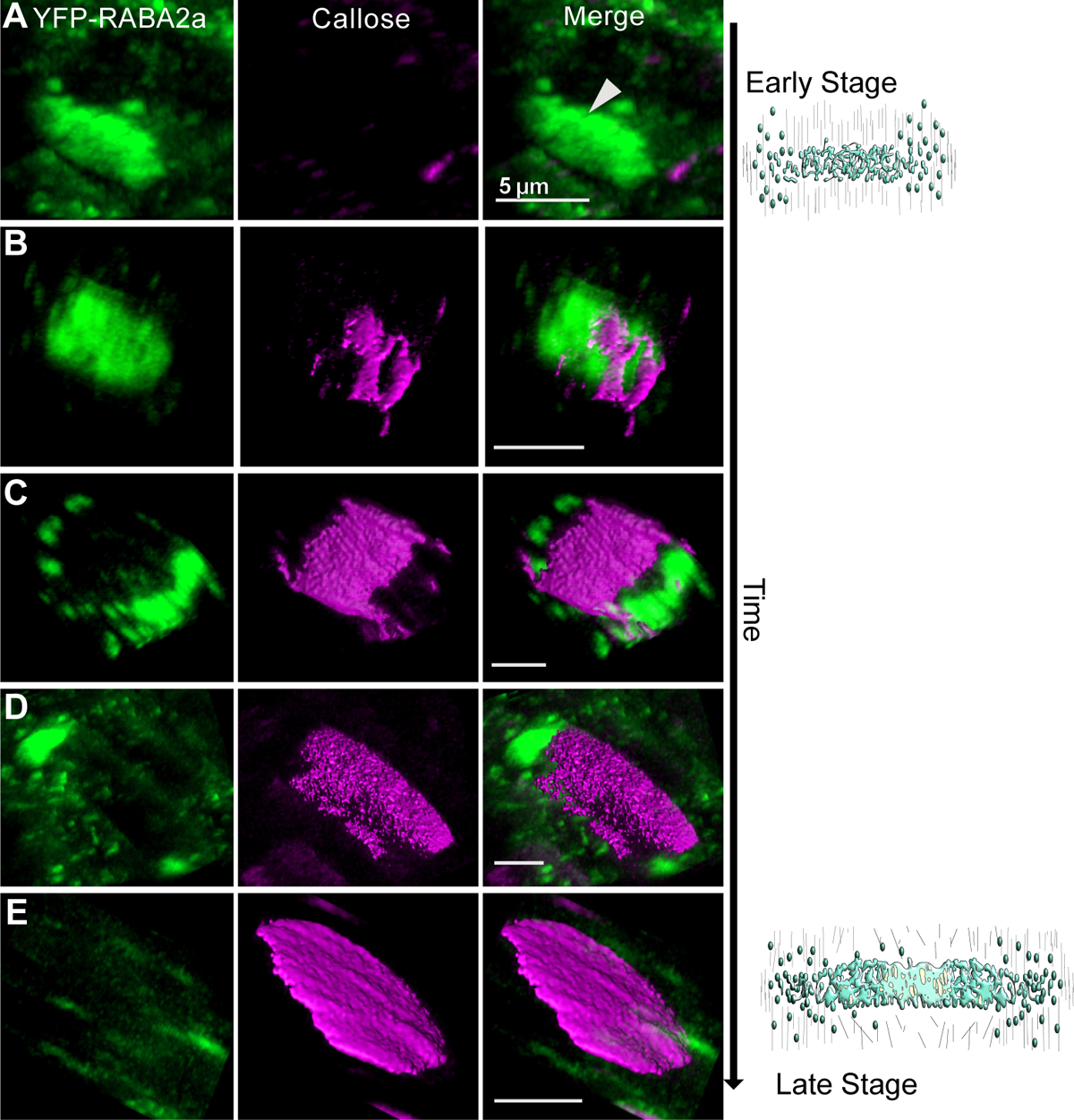
Progression of the cell plate in the presence of callose. **(A-E)** Cell plate progression in the presence of callose. **(A)** shows an early-stage cell plate before the accumulation of callose. At this stage, only vesicle accumulation by YFP-RABA2a (green) makes up the cell plate. **(B-D)** Later-stage cell plates where callose accumulation stained with aniline blue fluorochrome (magenta) is detectable. Note the transient accumulation of callose in later stages, leading to the maturation of the cell plate during normal cytokinesis. **(B)** Initial callose deposition overlapping with YFP-RABA2a. **(C)** As the cell plate maturation continues and the YFP-RABA2a accumulation takes a “doughnut shape” pattern, callose deposition appears in the middle of the cell plate with minimal overlap with YFP-RABA2a at the leading edge. **(D)** Callose is present throughout the cell plate, while minimal YFP-RABA2a is shown at the discontinuous ring **(E)**. Mature cell plate predominately labelled by callose. Scale bar = 5µm.

## Discussion

Mitosis, being the fundamental process of life that drives growth and development, necessitates a deeper understanding of its intricate details and complexity. Thus, the ability to image cytokinesis at a spatiotemporal level is of immense value in advancing developmental biology. In plants, cytokinesis is unique as it involves the *de novo* formation of a cell plate that expands centrifugally, leading to the separation of two daughter cells. The dynamic nature of various cell plate development stages that occur simultaneously demands sensitive imaging techniques with minimal photobleaching to comprehensively capture the entire process.

Although electron tomography studies have contributed significantly to our current knowledge of cytokinesis cytokinesis (Samuels et al., 1995; Segui-Simarro et al., 2004), they lack the ability to provide consecutive time points in the same sample, limiting their capacity for robust statistical analysis. Moreover, conventional confocal microscopy, despite its excellent lateral, axial, and temporal resolution, is too damaging to conduct high-quality, extended time-lapse acquisitions.

Additionally, the limited observable volume inherent in confocal microscopy, requires a manually guided acquisition onto the cell plate after it first becomes visible severely limiting the ability to capture early events and the ability to scale up observations.

The development of LLS microscopy affords imaging at much faster rates, with less light exposure of the sample, and minimizes photobleaching and phototoxic effects, which extends the observation time while affording higher axial resolution and maintaining confocal microscopy-type lateral resolutions (Chen et al., 2014). In this context, LLSM allows the collection of previously unobtainable datasets to dissect mitosis. Taking advantage of the resolution, speed, and gentle illumination, LLSM has been used to dissect mitosis and endomembrane dynamics in a variety of systems, including single cells in culture, *C. elegans*, zebrafish (Chen et al., 2014; Aguet et al., 2016), and human cells (Sen et al., 2021) providing new insights into cell division and its regulatory processes. The plant research community is slowly embracing the benefits that light sheet microscopy and vertical imaging may offer (Vyplelová et al., 2017; Glanc et al., 2018; Ovečka et al., 2021). Despite these forays, the powerful potential of LLSM has not been explored in plants to understand mechanisms that regulate plant cytokinesis. A likely factor is due to the limited availability of instrumentation and the effort required to establish an imaging routine along with quantitative analysis.

Further, given that plant cell walls may and did historically introduce aberrations in super-resolution imaging methods (Chatterjee et al., 2018; Novák et al., 2018; Ovečka et al., 2021), the performance of the LLMS in imaging plant samples is understandably a potential concern. This study, however, illustrates the feasibility and power of LLSM for attaining hitherto unobtainable detailed spatiotemporal data over time periods of 30-40 mins on a whole organismal level and within biologically relevant contexts in plants. Imaging at ∼25-30 sec time intervals allowed a detailed view of cell plate transition stages of high enough quality to segment and quantify at a biologically relevant scale. High imaging frequency is particularly exciting and valuable when observing transient events that appear somewhat randomly and unpredictably, such as cell plate formation within the intact root tip. The ability to capture complete (temporally and spatially) sporadic events embedded deep within plant tissue with sufficient resolution and statistically measurable numbers to quantitatively describe a complex as process such as cytokinesis represents a significant step forward in methodology and an opportunity for experimentation.

### Cell plate volume behavior follows three distinct and consistent phases

Our robust cytokinetic vesicle volume analysis during the entire process of cytokinesis showed distinct behavioral patterns that allowed the identification of three phases based on volume growth patterns. We attempt to assign previously structurally described cell plate development stages (Samuels et al., 1995) to the identified phases:

A. Phase I is characterized by a positive volume accumulation rate that peaks at ∼ 7-10 minutes before reaching an inflection point, representing the initial cell plate biogenesis stage. Our analysis showed a diameter increase relative to 50-60% of the parental cell width during phase I. The vesicle fusion, tubulovesicular network (FVS, TVN), and some tubular network TN transitions (Samuels et al., 1995) are likely included in this phase that corresponds to the delivery and fusion of cytokinetic vesicles at the cell plate edge and membrane network formation at the core of the cell plate disk.
B. Phase II, transitioning through an easily identifiable inflection point, is characterized by a negative volume growth rate with rapid loss reaching a minimum within ∼ 15 minutes. We reason that during this phase, representing a ring phragmoplast stage, the center of the cell plate has transitioned to a fenestrated sheet (PFS) (Samuels et al., 1995), requiring massive membrane recycling (Segui-Simarro et al., 2004). The observed recycling is consistent with earlier predictions estimating a 75% membrane reduction during cell plate maturation during endosperm cellularization (Otegui et al., 2001). The cell plate during this phase reaches almost 70-80% of the final cell plate diameter.
C. Phase III, marked by the remaining period of cell plate development, is characterized by a minimal but stable positive volume growth rate. The overall return to net positive growth likely represents PFS structures (Samuels et al., 1995; Segui-Simarro et al., 2004) associated with a discontinuous phragmoplast with minimal cytokinetic vesicle delivery, leading to the final expansion and maturation of the cell plate until it fuses with the parental cell wall.

Because cell plate development encompasses several stages that occur simultaneously in each phase, recycling of excess membrane material may already begin during phase I; however, based on the vast vesicle delivery, the net rate is positive contrasting with phase II, dominated by membrane recycling and a net negative volume growth rate. Phase III requires minimal membrane addition but stabilization and maturation of the cell plate, thus reverting to a net positive volume growth rate. The minimal return to net positive increase could be due to the reorganization of the cell plate membrane at the edge, as it needs to combine with the parental plasma membrane carrying the targeted and necessary machinery and cargo necessary for joining with the parental cell wall.

Earlier models based on cell plate diameter suggest a gradual decrease in cell plate expansion rates (Higaki et al., 2008) or a biphasic pattern (van Oostende-Triplet et al., 2017) as the cell plate expands towards the parental wall. The van Oostende-Triplet et al. model identifies a Primary Centrifugal Growth (PCG) phase in BY-2 cells with an average diameter growth rate of 1.44 ± 0.44 µm min^−1^ and Secondary Centrifugal Growth (SCG) phase with a rate of 0.35 ± 0.13 µm min^−1^ along with similar estimates in Arabidopsis (PCG ∼0.87 ± 0.26 µm min^−1^ and SCG ∼0.25 ± 0.09 µm min^−1^) (van Oostende-Triplet et al., 2017). Analysis of our diameter expansion data, as guided by the RABA2a marker, showed an initial high growth rate 4.09 ± 0.26 µm min^−1^ followed by a gradual decrease which can be grouped in two bins that average 1.1 ± 0.05 µm min^−1^ and 0.59 ± 0.04 µm min^−1^ (n=18) (**Fig. S4A**), consistent with the earlier calculated rates described above (van Oostende-Triplet et al., 2017). The extremely rapid expansion rate within the first 2.5 minutes of cell plate development (**Fig. S4B, C**) aligns with cell plate biogenesis during vesicle delivery. It corresponds to the Initial Plate Assembly (IPA) stage by van Oostende-Triplet et al., 2017, for which expansion rates were not detectable. Taken together, this is a clear example of LLSM application in enabling the recording of cell plate biogenesis and the calculation of the corresponding rates, as it can capture early events prior to well -defined cell plate appearance in the field of view.

It is apparent in comparing different models that understanding cell plate development based on volumetric growth provides more straightforward cutoff points for phase transitions and complements cell plate diameter analysis. Proof of this idea is demonstrated in this study with the volume accumulation rates of the easily trackable cytokinesis marker. This analysis allows the prediction of quantitative behaviors during cell plate development and their interrogation to understand the contribution of different components. For example, models of phragmoplast expansion (Higaki et al., 2008) can be reviewed with the analysis proposed here to evaluate the contributions of the array that helps build the cell plate. The easily identifiable inflection points between phases mark critical points during the transition of the membrane network and cell plate expansion that requires the onset of specific contributing factors. We propose that during the transition from phase I to II, clathrin-mediated recycling is enhanced, and that polysaccharide deposition and assembly are dominant during the transition from phase II to III. Future studies can explore the method and proposed model to interrogate different markers such as clathrin, dynamin, and SNARE proteins involved in different aspects of cell plate development. It is plausible that the volumetric-based transitions will vary based on the marker used e.g., SNARE versus clathrin (Bednarek and Backues, 2010; El Kasmi et al., 2013; Jurgens et al., 2015; Dahhan and Bednarek, 2022; Park et al., 2023), which will help build more comprehensive models.

### Callose is necessary during cell plate expansion for the transition beyond phase I

An example of how the current analysis and proposed model can provide insights into cell plate development is the contribution of callose, a polysaccharide transiently deposited during cell plate development. The specific stage for which callose plays a critical role has been long debated, with views arguing on either the transition from a TN to PFS or the cell plate insertion to the parental cell wall (Samuels et al., 1995; Thiele et al., 2009). Different methods to dissect its role, direct detection of callose with antibodies against the polysaccharide with EM (Samuels et al., 1995) versus confocal 2D staining and mutant characterization (Thiele et al., 2009) might contribute to these diverging hypotheses. The developed approach and the derived model here, in combination with pharmacological treatment, allow the detailed dissection of each stage, providing insights into the biological role of the polymer. Upon inhibition of callose with ES7, phase I is prolonged. An irregular pattern is seen following the initial phase, with no distinguishable phases II and III. This clearly illustrates that polysaccharide is essential during the transition beyond phase I.

A modeling approach was previously implemented to understand better the cell plate stage transition from a vesicular network to a fenestrated sheet and mature cell plate. The model predicts that the onset of a two-dimensional spreading/stabilizing force, coupled with a concurrent loss of spontaneous curvature, is necessary for cell plate expansion (Jawaid et al., 2022). Biophysical modeling highlights the need for a spreading force during the transition of the membrane network (TN) to a fenestrated sheet (PSF) (Jawaid et al., 2022), which overlaps with the phase transition (Ph I to Ph II) in our study. Directly detecting callose via fluorescent staining corresponds to these phases and validates the prediction of the biophysical model. Callose, due to its amorphous structure,can lead to alterations in the physical and mechanical properties at the deposition site (Piršelová and Matušíková, 2012; Zhang et al., 2021; Usak et al., 2023). The polysaccharide has the ability to enhance rigidity while maintaining flexibility and reduce permeability to various compounds across a range of functions (Yim and Bradford, 1998; Parre and Geitmann, 2005; Vaten et al., 2011). Additionally, callose’s unique composition compared to other polysaccharides enables controlled degradation when no longer necessary, undelying its significance as a compound that can operate both spatially and temporally (Samuels et al., 1995; Usak et al., 2023).

### LLSM along with 4D volume-based phase transition is robust in cytokinesis dissection

The quantitative prediction of phase transition, with unbiased characterization, provides an advantage over descriptive interpretations of fragmented data sets. Further, it establishes a roadmap into which other components can be incorporated. Structural cell wall proteins, such as extensins (Cannon et al., 2008), wall matrix polysaccharides (Moore and Staehelin, 1988), cellulose (Miart et al., 2014), and other forms of linear glucose can be interrogated with the same analysis methodology. Cellulose, the load-bearing cell wall polysaccharide, is a prime candidate for this analytical approach, especially in combination with conditional genetic mutations or pharmacological inhibition of cellulose synthases (Chen et al., 2018). Interestingly, a five-day inhibition of cellulose synthase (CESA) activity with isoxaben causes a reduction of cell elongation but does not have a prominent effect on cell plate biogenesis or expansion (Jawaid et al., 2022), which suggests a role of cellulose in the formed cell wall instead of cell plate expansion. A quantitative analysis during cell plate phase transition can help dissect the contribution of cellulose or other linear β-1,4-glucans. Beyond cellulose synthase, cellulose synthase-like proteins (CSLs), such as CSLD3, produce linear glucan polymers (Yang et al., 2020). Cellulose synthase D5 (CSLD5), a cytokinesis-specific protein (Gu et al., 2016), produces a β-1,4-glucan polysaccharide, similar to that of CSLD3 (Yang et al., 2020). Application of our quantitative analysis in conditional mutations of these proteins can pin down the specific contribution to cell plate development and indicate associations with other polysaccharides or cell wall components. Such a study can address the potential interaction of CSLD5 products with callose in creating a scaffold (Abou-Saleh et al., 2018) for the establishment of the spreading force necessary for cell plate maturation.

While our study was centered on callose, the described model of cytokinetic vesicle behavior can be interrogated for an array of contributions to cell plate development (Smertenko, 2018; Cheng and Bezanilla, 2021; Dahhan and Bednarek, 2022; Gu and Rasmussen, 2022; Sinclair et al., 2022; Lebecq et al., 2023; Nan et al., 2023; Park et al., 2023). For example, pharmacological inhibition of secretory traffic, cytoskeleton dynamics at the phragmoplast, and its interaction with vesicle delivery can dissect the contribution of each element. In a recent study, the application of the Small Molecular Inhibitor Formin Homology 2 (SMIFH2) revealed the function of formins in several aspects of cytokinesis, including cell plate membrane organization, as well as microtubule polymerization and nucleating F-actin at the cell plate (Zhang et al., 2021). The application of LLSM and the methodology developed here can aid in further dissecting the role of formins in cell plate development and phragmoplast organization. The pharmacological inhibition of actin or microtubule polymerization has been extensively used to study phragmoplast expansion and recently shown for cell plate development using a radial expansion model (van Oostende-Triplet et al., 2017). Extending these studies to the 4D analysis based on rates of volume accumulation described here provides extra depth, allowing for direct correlations of the contributions of cytoskeleton dynamics, secretory traffic, and protein synthesis in cell plate biogenesis and expansion.

## Conclusion

The development of a comprehensive 4D image acquisition and processing pipeline using LLSM represents a significant step forward in both utilizing cutting-edge microscopy tools and addressing long-standing questions about cytokinesis spatiotemporal dynamics. Despite the advantages, today there are still limits for LLSM based imaging of plant seedlings. The current approach requires genetically encoded fluorophores with high photostability, thus not all markers are ideally suited. Multiplex imaging of different fluorophores may be impacted by cross talk and requires multichannel camera systems. Another major challenge is the substantial data size generated during the image analysis and archiving pipeline. For instance, the current study produced around 40TB of data, which poses significant resource demands on IT infrastructure for processing and storage. However, it is important to highlight a distinctive feature of this experimental design—the ability to conduct prolonged imaging over a large field of view. This capability allows researchers to capture the unpredictable appearance of cell plate formation from its initial stages until completion and observe other unique imaging characteristics offered by LLSM.

The developed 4D imaging pipeline using LLSM, along with the tools for processing the acquired images are documented and are available through plugins in ImageJ and MATLAB. The segmentation pipeline employed in Bitplane’s Imaris is also documented for easy adaptation (**Fig. 2**). The cell plate development model can be applied using different modalities beyond LLSM to dissect plate cytokinesis. We hope that this study inspires the community to adopt the methodology of exploring LLSM in plants, take advantage of the image analysis pipeline tools developed, and refine the model of cell plate development and its factors contributing to each phase.

More importantly our study was able to showcase first-of-its-kind data of the complete process of cell plate development with minimal photobleaching, higher axial resolution, and faster imaging rates compared to traditional confocal microscopy. The study revealed three distinct developmental phases of cell plate growth, from rapid vesicle accumulation to subsequent volume reduction and cell plate expansion and the critical role of callose in phase transition. The use of the chemical inhibitor ES7 in combination with LLSM provided quantitative insights into the timing and role of callose in stabilizing the cell plate during expansion and maturation. Inhibition of callose deposition led to impaired cell plate expansion, resulting in fragmented and aberrant membrane accumulation patterns. The findings of this study have significant implications for understanding the spatiotemporal dynamics of cell plate development and the role of callose along with other components in this process.

We see this pioneering effort in quantitative/modeling dissection of cell plate development that will help understand the array of factors controlling plant cytokinesis. Taken together, this research contributes to the broader understanding of plant cell biology and opens new avenues for further investigations into the molecular mechanisms underlying cell plate assembly and expansion.

## Abbreviation

3D: three-Dimensional

4D: four-dimensional

AIC: advanced imaging center

CSLs: cellulose synthase-like proteins

ES7: endosiden 7

FVS: fusion of vesicle stage

GPU: graphics processing unit

GSL8: glucan synthase like 8 / callose synthase

LLSM: lattice light sheet microscopy

MS: Murashige and Skoog

PFS: planar fenestrated sheet

Ph: Phase

PSF: point spread function

RANSAC: Random sample consensus

ROI: Region of Interest

SMIFH2: Small Molecular Inhibitor Formin Homology 2

TEM: transmission electron microscopy

TN: tubular network

TVN: transition to a tubular-vesicular network

## Acknowledgments

This manuscript is dedicated in memory of Prof. Andrew Staehelin who’s work and contribution to the field inspired us in this study. We want to thank all the members of AIC and the Drakakaki lab for their input and Dr. Destiny J. Davis for her critical reading of the manuscript. We would like to acknowledge the use of Zeiss 980 confocal microscope made available through the NIH grant S10OD026702

## Author contributions

Conceptualization: RMS, TW, GD; Methodology: RMS, MW, MZJ, TW, GD; Formal analysis: RMS, MZJ, DC, JH, TW; Investigation: RMS, MW; Resources: RMS, KMD, TW, GD; Data curation: RMS, MW, GD; Writing - original draft: RMS, MZJ, DC, TW, GD; Writing - review & editing: RMS, MW, MZJ, TW, GD; Validation: RMS, MZI; Visualization: RMS, JA, KMD, JH, TW, GD; Supervision: GD; Software: RMS, MW, MZJ, BR, EW, KMD, DC, JH; Project administration: GD; Funding acquisition: DC, GD; Design research, Performed research analyze data, provided recourses, Wrote manuscript, edited manuscript

## Supplemental Figures

**Fig. S1.** Statistical comparison of volume accumulation. YFP-RABA2a normalized volume growth rates, averaged within predefined groups and shown with 95% confidence intervals. **(A-C)**. Bins 1-3 represent the rates corresponding to normalized volumes increasing from 0-0.33 µm^3^, 0.33-0.66 µm^3^, and 0.66-1 µm^3^, respectively. All bins correspond to the first derivative rates of volume growth. * Indicates P < 0.005 **(D-E)** Bins 4,5 represent the rates corresponding to volumes decreasing from 1-0.66 µm^3^ and 0.66-0.33 µm^3^, respectively; bin 6 data are not shown due to the bin spanning over the region in which the end of collection has limited control over noise. All bins correspond to the first derivative rates of volume growth. * Indicates P < 0.005 **F)** Schematic representation of volume growth grouped in bins for further processing as shown in **A-E**.

**Fig. S2.** Max accumulation of volume and surface area across treatments. YFP-RABA2a maximum volume and surface area comparisons averaged across all cell plates analyzed and shown with 95% confidence intervals. The lack of statistical significance supports the hypothesis that ES7 does not impact vesicle delivery but the ability to mature the cell plate into a proper cross wall.

**Fig. S3.** Individual cell plate rates of expansion. Expansion rates of individual cell plates based on their radial growth. **(A, C)**. Each line indicates cell plate diameter for individually segmented cell plates represented in Fig. 3 and Fig. 5, respectively, to untreated **(A)** or ES7 treated conditions **(C)**. Normalized diameters to their final cross-wall length of the developing cell plates were fitted to a second-degree polynomial distribution for both untreated **(A)** and ES7 treated samples **(C)**. First-degree derivatives for untreated **(B)** and ES7 treated **(D)** samples are shown. E) Polynomial fitted cell plate diameters were grouped into 20 intervals (“bins”) which were subjected to statistical analysis comparing control versus ES7 treated cell plates as shown in Fig. 6D.

**Fig. S4.** Comparison of expansion trends based on diameter or volume. Each cell plate’s radial growth rate was measured during development and binned by time, based on growth rate values **(A)**. The first bin is defined by a rapid increase during 0-up to 2.42 ± 0.05 min with an average growth rate of 4.09 ± 0.26 µm min^−1^. Bin two takes place between 2.42 ± 0.05 min to 15.07 ± 0.4 min where the radial growth rate gradually decreases to 1.1 ± 0.05 µm min^−1^, and finally bin three within the remaining time up to 24.82 ± 1.43 min comprised of radial rates averaging 0.59 ± 0.04 µm min^−1^ (n=18) (see Fig. 6). **B)** shows the radial expansion rates, for each bin, with an overall downward trend. **C)** Further the radial expansion rate groups (bins) were overlaid to the volume growth rates of the representative cell plate highlighted in Fig. 3 **A-C.**

**Video. S V1**. 4D rendering of YFP-RABA2a cell pate accumulation with surface segmentation. Images were acquired with LLSM. Scale = 20 µM

**Video. S V2.** 4D rendering of YFP-RABA2a cell pate accumulation with surface segmentation under 2 hour 50 µM Endosidin 7 treatment displaying relatively no expansion. Images were acquired with LLSM. Scale = 20 µM

**Video. S V3.** 4D rendering of YFP-RABA2a cell pate accumulation with surface segmentation under 2 hour 50 µM Endosidin 7 treatment displaying minimal relative expansion. Images were acquired with LLSM. Scale = 20 µM

