## Supplementary material for "Four-dimensional quantitative analysis of cell plate development using lattice light sheet microscopy identifies robust transition points between growth phases": Sup1

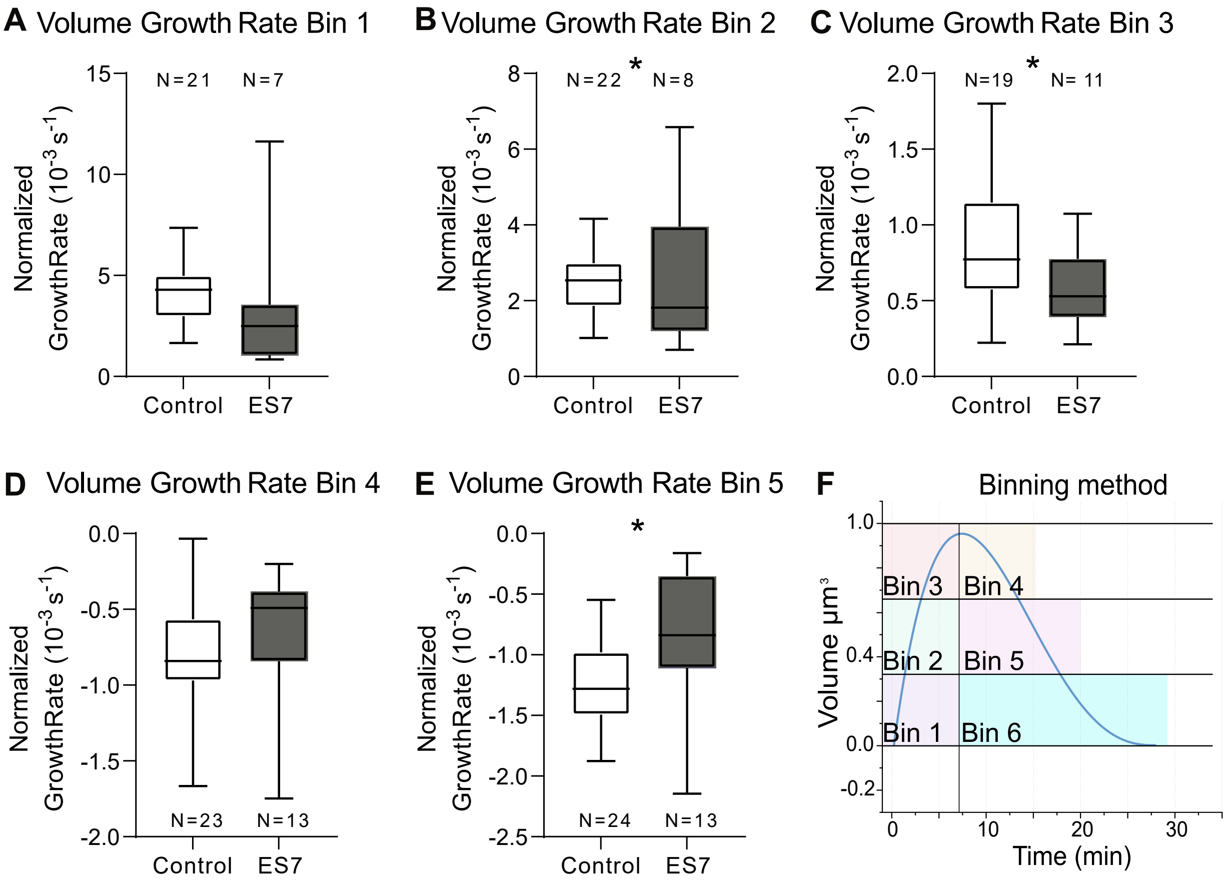


**Fig. S1.** Statistical comparison of volume accumulation.

YFP-RABA2a normalized volume growth rates, averaged within predefined groups and shown with 95% confidence intervals.

**(A-C)**. Bins 1-3 represent the rates corresponding to normalized volumes increasing from 0-0.33 μm$^3$, 0.33-0.66 μm$^3$, and 0.66-1 μm$^3$, respectively. All bins correspond to the first derivative rates of volume growth. * Indicates P < 0.005

**(D- E)** Bins 4,5 represent the rates corresponding to volumes decreasing from 1-0.66 μm$^3$ and 0.66-0.33 μm$^3$, respectively; bin 6 data are not shown due to the bin spanning over the region in which the end of collection has limited control over noise. All bins correspond to the first derivative rates of volume growth. * Indicates P < 0.005

**F)** Schematic representation of volume growth grouped in bins for further processing as shown in **A-E**.
