## Supplementary material for "Four-dimensional quantitative analysis of cell plate development using lattice light sheet microscopy identifies robust transition points between growth phases": Sup2

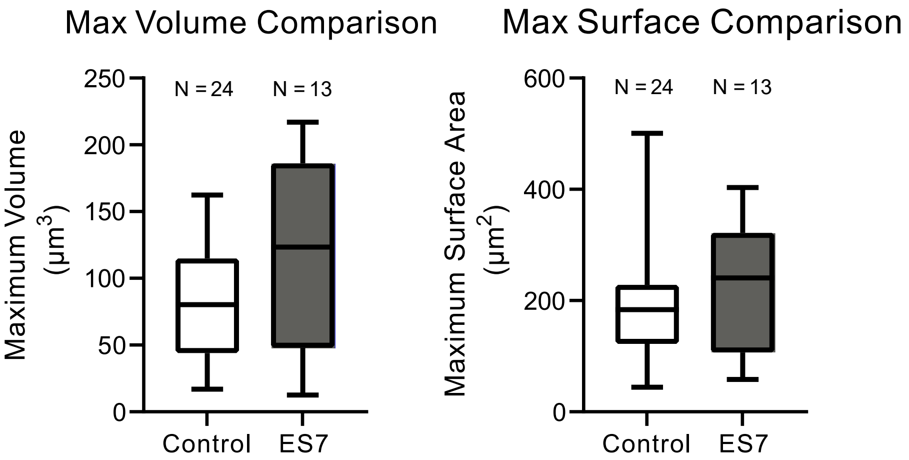


**Fig. S2.** Max accumulation of volume and surface area across treatments.

YFP-RABA2a maximum volume and surface area comparisons averaged across all cell plates analyzed and shown with 95% confidence intervals. The lack of statistical significance supports the hypothesis that ES7 does not impact vesicle delivery but the ability to mature the cell plate into a proper cross wall.
