## Supplementary material for "Four-dimensional quantitative analysis of cell plate development using lattice light sheet microscopy identifies robust transition points between growth phases": Sup3

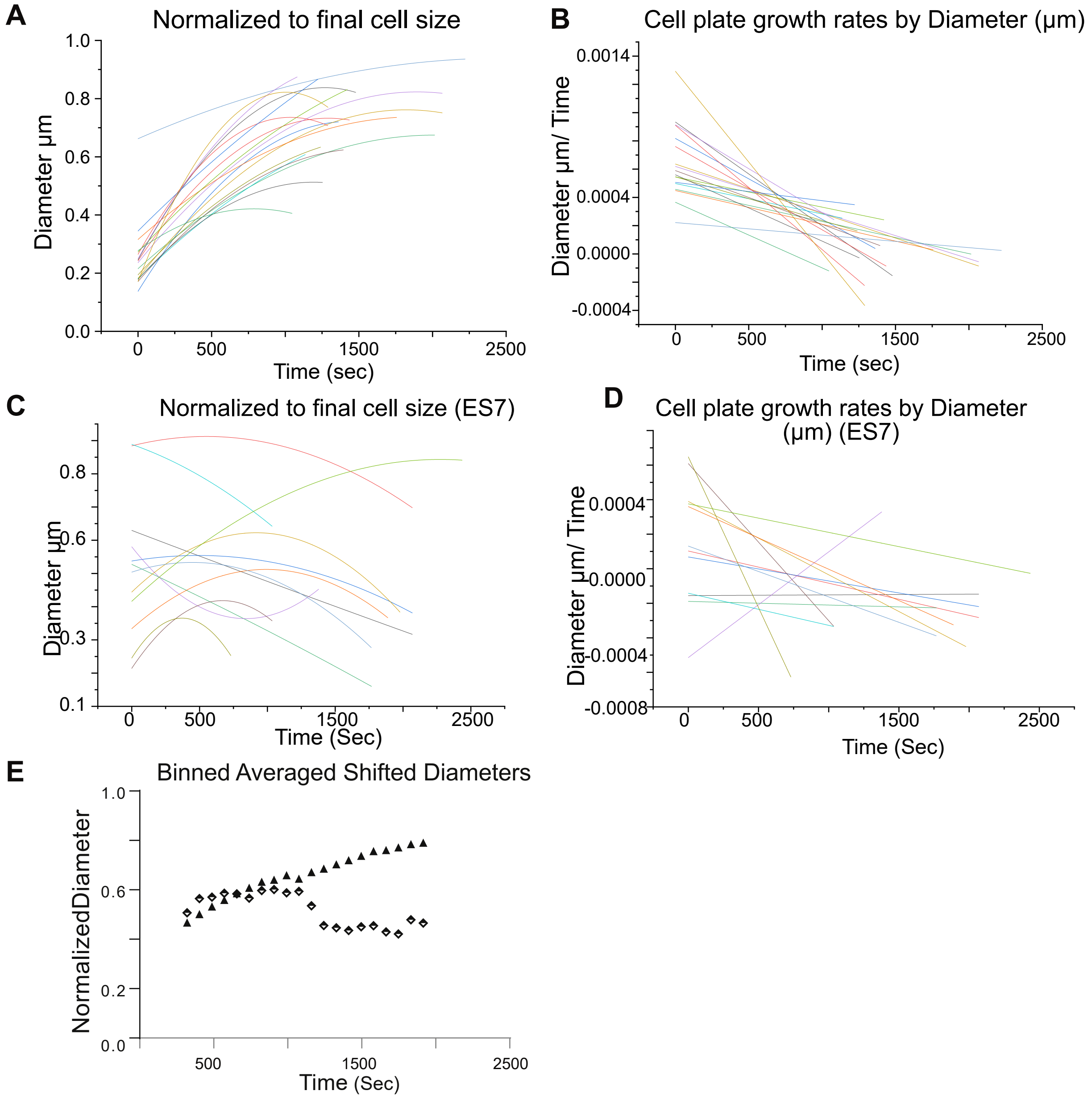


**Fig. S3.** Individual cell plate rates of expansion.

Expansion rates of individual cell plates based on their radial growth. **(A, C)**. Each line indicates cell plate diameter for individually segmented cell plates represented in Figure 3 and Figure 5, respectively, to untreated **(A)** or ES7 treated conditions **(C)**. Normalized diameters to their final cross-wall length of the developing cell plates were fitted to a second-degree polynomial distribution for both untreated **(A)** and ES7 treated samples **(C)**. First-degree derivatives for untreated **(B)** and ES7 treated **(D)** samples are shown. E) Polynomial fitted cell plate diameters were grouped into 20 intervals ("bins") which were subjected to statistical analysis comparing control versus ES7 treated cell plates as shown in Figure **6D.** non treated n=24 Treated n=18
