## Supplementary material for "Four-dimensional quantitative analysis of cell plate development using lattice light sheet microscopy identifies robust transition points between growth phases": Sup4

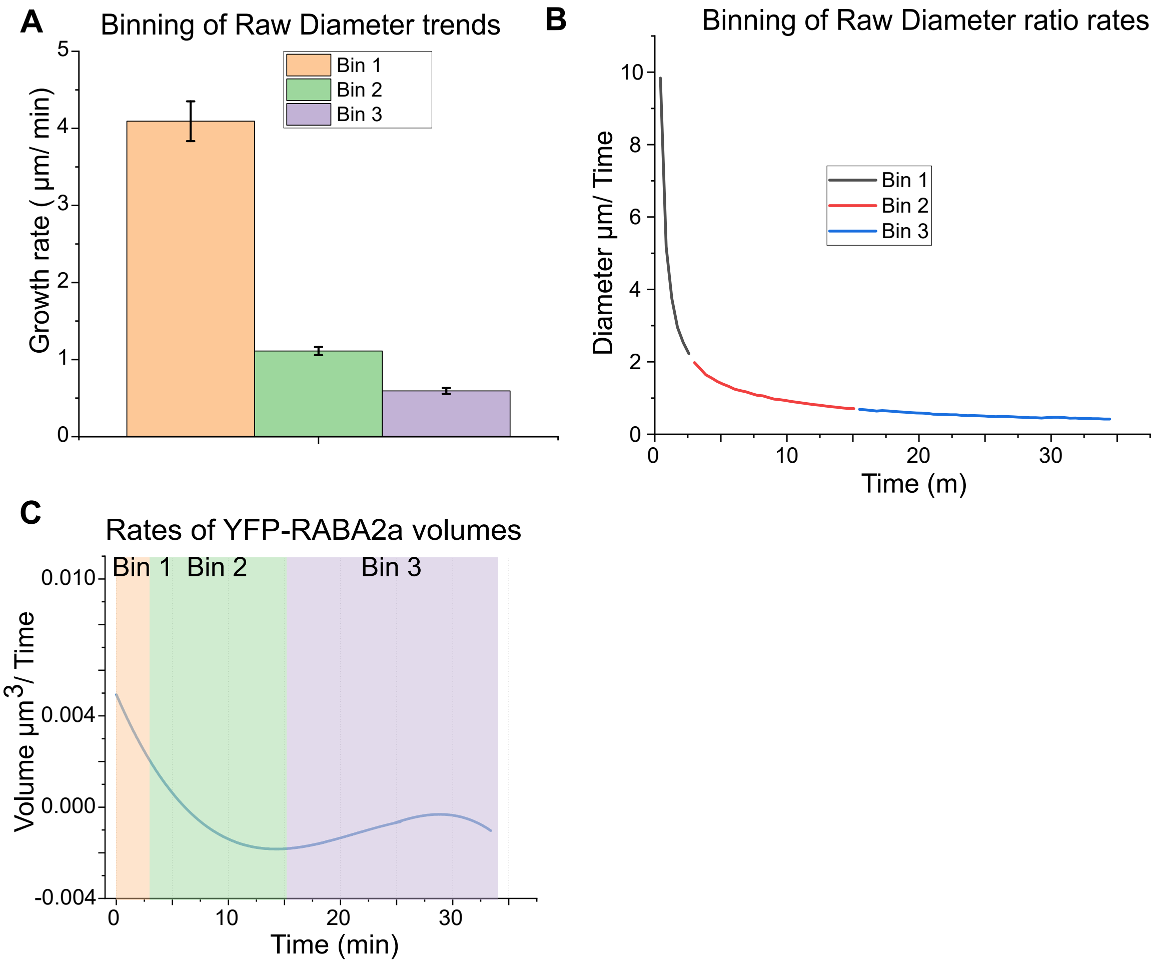


**Fig.S4.** Comparison of expansion trends based on diameter or volume.

Each cell plate's radial growth rate was measured during development and binned by time, based on growth rate values **(A)**. The first bin is defined by a rapid increase during 0-up to 2.42 ± 0.05 min with an average growth rate of 4.09 ± 0.26 µm $min^{-1}$. Bin two takes place between 2.42 ± 0.05 min to 15.07 ± 0.4 min where the radial growth rate gradually decreases to 1.1 ± 0.05 µm $min^{-1}$, and finally bin three within the remaining time up to 24.82 ± 1.43 min comprised of radial rates averaging 0.59 ± 0.04 µm $min^{-1}$ (n=18) (see **Figure 6**).

**B)** shows the radial expansion rates, for each bin, with an overall downward trend. **C)** Further the radial expansion rate groups (bins) were overlaid to the volume growth rates of the representative cell plate highlighted in **Figure 3 A-C.**
